# Polyhydroxybutyrate production in freshwater SAR11 (LD12)

**DOI:** 10.1101/2024.11.08.622676

**Authors:** Brittany D. Bennett, David A.O. Meier, V. Celeste Lanclos, Hasti Asrari, John D. Coates, J. Cameron Thrash

**Affiliations:** Marine and Environmental Biology, University of Southern California, Los Angeles, USA; Department of Plant and Microbial Biology, University of California, Berkeley, Berkeley, USA

## Abstract

SAR11 bacteria (order *Pelagibacterales*) are oligotrophs and often the most abundant bacterioplankton in aquatic environments. A subset of sequenced SAR11 genomes, predominantly in the brackish and freshwater SAR11 subclades, contain homologs of *pha* genes, which in other organisms confer the ability to store carbon and energy via polyhydroxyalkanoate (PHA) polymers. Here, we investigated the relevance of PHA production to SAR11 biology. Phylogenetics showed that Pha proteins occurred on a long branch and provided evidence for origin at the common ancestor of the brackish IIIa and freshwater LD12 subclades, followed by horizontal transfer within SAR11. Using the LD12 representative “*Candidatus* Fonsibacter ubiquis” strain LSUCC0530, we found that a large minority of LSUCC0530 cells contained a single Nile red-staining granule, confirmed that the cells produced polyhydroxybutyrate (PHB), and estimated the total PHB content in the cells. We heterologously expressed the LSUCC0530 *phaCAB* locus in *Escherichia coli*, finding it to be functional and the likely origin of the PHB. We also determined that, irrespective of changes to carbon, nitrogen, and phosphorus concentrations, a similar fraction of LSUCC0530 cells generated PHB granules and expression of the *phaCAB* locus remained constant. We suggest that PHB synthesis in LSUCC0530 may be constitutively active due to the slow growth dynamics and minimal regulation that characterize SAR11 bacteria. This is the first characterization of polymer storage in SAR11, providing new insights into the likely fitness advantage for cells harboring this metabolism.

## Introduction

Alphaproteobacterial strains within the order *Pelagibacterales*, colloquially the SAR11 clade, are ubiquitous, highly abundant, slow-growing members of the bacterioplankton in aquatic ecosystems ranging from marine to freshwater (1–4). Although the majority of research on SAR11 has focused on marine oligotrophs that frequently dominate nutrient-poor offshore surface communities (3,5,6), including the well-described SAR11 subclade Ia strain “*Candidatus* Pelagibacter ubique” HTCC1062 (2), SAR11 also comprises groups that are adapted for brackish (subclade IIIa) (7) and fresh water (subclade IIIb, also called LD12) (8,9), which often make up the largest fraction of low salinity bacterioplankton (10).

Regardless of environmental habitat, all SAR11 bacteria have highly streamlined genomes with limited metabolic pathways and few annotated regulatory genes (9,11,12), resulting in a high degree of constitutive gene expression (12–14), making them both particularly fit for low-nutrient conditions and largely unable to respond transcriptionally or physiologically to changes in their environmental conditions (4,9,11,12). SAR11 genomes also share large percentages of their genes even across subclades, resulting in similar overall metabolic profiles despite hundreds of millions of years of divergent evolution (9,15). Nevertheless, a distinguishing genomic feature that may provide a means of niche adaptation in some SAR11 lineages is the potential for carbon polymer storage (7,16,17). Homologs of polyhydroxyalkanoate (PHA) synthesis genes occur in multiple SAR11 isolate and metagenome-assembled genomes from both brackish and freshwater environments, predominantly in the IIIa and LD12 subclades (7,16,17). However, the functions and possible fitness benefits this pathway might confer on the few SAR11 groups that harbor it still need investigation.

PHA production is a carbon and energy storage mechanism widespread throughout the Bacteria (18). In fact, some PHA-producing microbes can store up to ∼70% of their dry mass as PHA (19–21). Depending on the *pha* genes present in an organism’s genome and the available carbon source(s), PHA polymers composed of monomers with varying side chain lengths and structures may be produced (22), though the most well-studied PHA is polyhydroxybutyrate (PHB). A common PHA production pathway requires three core enzymes: acetyl-CoA acetyltransferase PhaA (in some organisms designated PhbA), acetoacetyl-CoA reductase PhaB (or PhbB), and polyhydroxyalkanoate synthase PhaC (or PhbC) (23,24). Together these enzymes form a three-step pathway in which (in organisms producing PHB) two acetyl-CoA molecules are synthesized into acetoacetyl-CoA, which is reduced to the monomer unit 3-hydroxybutyryl-CoA, and then these monomers are polymerized into PHB. Dozens of alternative monomer units, such as 3-hydroxyvalerate, 3-hydroxyoctanoate, and 3-hydroxydecanoate, have been found incorporated into PHA polymers (25).

Regulation of PHA production is multilayered and can occur at the transcriptional, translational, and/or post-translational levels, though the mechanisms vary by organism (reviewed in (26,27)). Most regulatory mechanisms discovered thus far appear to affect the amount of PHA accumulated in the cell, whereas the monomer composition is usually determined by the carbon source(s) provided to the organism (28). Generally, a combination of nutrient limitation, particularly of nitrogen and/or phosphate, and carbon abundance will induce PHA storage in organisms that have this capacity (29,30). Carbon may be shunted toward PHA formation when other nutrients have been exhausted due to allosteric regulation of PhaC and/or PhaA by metabolic pathway intermediates (31–33). Additionally, in various organisms there is a range of upstream mechanisms that regulate the *pha* locus in response to environmental conditions (reviewed in (27)). Given the low number of regulatory elements encoded in SAR11 genomes and their oligotrophic adaptations, it was unclear to us what, if any, regulatory mechanism(s) governing PHA production might exist in this clade. Furthermore, the type of PHA produced and the amount stored per cell are important data for understanding how SAR11 organisms use this metabolism.

In this work, we present evidence that carbon storage in the form of PHA polymers may confer a fitness benefit to the cultured LD12 representative “*Candidatus* Fonsibacter ubiquis” strain LSUCC0530. Previously, a putative *pha* locus was identified in the LSUCC0530 genome (7). Here we use genetic, phenotypic, and chemical analyses to demonstrate that this locus is functional and that LSUCC0530 produces PHA granules under all tested growth conditions. We postulate that LSUCC0530 synthesizes PHA regardless of environmental nutrient levels, and that this stored carbon may aid the bacterium during starvation periods. Furthermore, the phylogenetic similarity of LD12 PHA production genes to those in other SAR11 strains suggests a common strategy that may have been horizontally transferred within the clade.

## Materials and Methods

### Bacterial strains and growth conditions

Bacterial strains and plasmids used in this study are listed in Table 1. We grew LSUCC0530 in polycarbonate flasks at room temperature in either JW5 (Table S1) (34) or CCM5PK (Table S1), and HTCC1062 in polycarbonate flasks at room temperature in AMS1 supplemented with 10 µM L-methionine, 50 µM L-glycine, and µM sodium pyruvate (35). SAR11 strains were grown without shaking unless otherwise noted. We grew *Escherichia coli* at 37°C in Lennox Luria Broth (Sigma-Aldrich, St. Louis, MO) supplemented with 50 μg/ml kanamycin (Sigma-Aldrich), 300 µM diaminopimelic acid (Sigma-Aldrich), and/or 200 mM isopropyl ß-D-1-thiogalactopyranoside (AmBeed, Arlington Heights, IL) where applicable; liquid cultures were shaken at 180 rpm. We grew *Cupriavidus necator* H16 at 30°C in nutrient broth (BD Difco, Franklin Lakes, NJ) supplemented with 0.2% (wt/vol) sodium gluconate when necessary to induce PHA production; liquid cultures were shaken at 180 rpm.

**Table 1.**
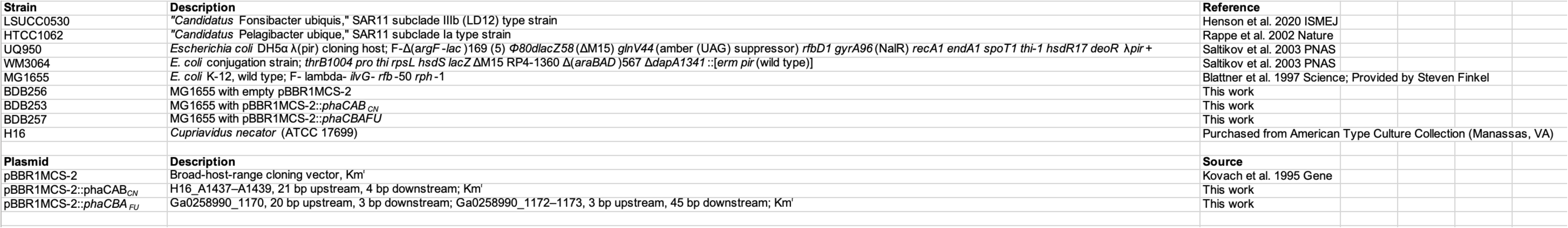
Bacterial strains and plasmids used in this study.

### Growth curves

We inoculated cryostocks of SAR11 strains into triplicate 25 ml starter cultures of either JW5 (LSUCC0530) or AMS1 (HTCC1062) and grew them to late exponential phase. We generated growth curves by inoculating each starter culture to a cell density of ∼2500 cells/ml into 25 ml of either JW5 (LSUCC0530) or AMS1 (HTCC1062), with or without organic carbon sources as noted. We monitored growth by periodically staining aliquots with 1x SYBR green I (Lonza, Basel) and enumerating them on an Accuri C6 Plus flow cytometer (BD Biosciences) (ex 488 nm, em 533/30). We calculated growth rates in Prism 5 by first fitting a nonlinear regression (Y=Y_0_*exp(k*X), where Y=cells/ml, X=time, and k=rate constant) to the exponential phase of the growth curve, then calculating doubling time=ln(2)/k.

### Transmission electron microscopy

We grew LSUCC0530 cells in JW5 to late exponential phase, processed them as described previously (7), and took images on a JEM-1400 transmission electron microscope. PHB content was estimated by measuring the volume of each inclusion body in Fiji ver. 2.15.0 running ImageJ ver. 1.54f (36) and multiplying by the density of amorphous PHB (37).

### Fluorescence microscopy

We prepared LSUCC0530 cultures as follows: 2 ml of culture was centrifuged for 30 min at 17,500 x g. The top 1.92 ml of supernatant was carefully pipetted off and discarded, and 500 µL 35% EtOH in PBS (per L: 8 g NaCl, 201 mg KCl, 1.42 g Na2HPO4, 245 mg KH2PO4, pH 7.4) was added to the cell pellet. Cells were vortexed vigorously and incubated 15 min at room temperature. Cells were centrifuged again for 30 min at 17,500 x g, after which the top 500 ul of supernatant was carefully pipetted off and discarded. 200 µl PBS was added to each cell pellet, and pellets were vortexed vigorously. 0.6 µl 1 mg/ml DAPI (Thermo Scientific, Waltham, MA) and 0.6 µl 10 mg/ml Nile red (Sigma-Aldrich) in 100% DMSO were added to the cell suspension, and cells were mixed gently by inversion and incubated in the dark for 30 min. 6 µl stained cells were mounted onto a slide affixed with a 1% agarose pad (38), topped with a #1 coverslip followed by a drop of Type B immersion oil (Cargille, Cedar Grove, NJ), and imaged on an Eclipse Ti epifluorescence microscope (Nikon Instruments, Melville, NY) equipped with a Plan Apo 100x/1.40 oil immersion objective, X-Cite 110LED light source (Excelitas Technologies, Waltham, MA), and DAPI (ex 360-380 nm, em 435-485 nm) and red-shifted TRITC bandpass (ex 528-553, em 590-650) filter cubes.

We prepared *E. coli* cultures as follows: 100 µl liquid overnight culture was centrifuged for 1 min at 8000 x g. The supernatant was poured off and discarded, and the cell pellet was resuspended in 500 µl PBS and then centrifuged for 1 min at 8000 x g. The supernatant was discarded, and cells were resuspended in 500 µl 35% EtOH in PBS and then incubated at room temperature for 15 min. Cells were centrifuged for 1 min at 8000 x g, and the supernatant was discarded. The cell pellet was resuspended in 500 µl PBS, and 0.5 µl 1 mg/ml DAPI and 1.5 µl 1 mg/ml Nile red in 100% DMSO were added. Cells were mixed gently by inversion and incubated in the dark for 30 min before mounting onto slides and imaging as above.

### Phylogeny of Pha proteins

We queried the PhaC, PhaA, and PhaB protein sequences from LSUCC0530 and the PhaC1, PhaA, and PhaB1 protein sequences from *C. necator* H16 twice in PSI-BLAST (39) against the RefSeq Select database: once against the full database, and once limiting the search to the SAR11 cluster, using as many iterations as necessary to reach convergence. We selected the top 250 proteins returned for each search, minus “Multispecies” hits, for inclusion in the analysis. To fill in the SAR11 section of the tree and represent a broad range of strains, collection sites, and query coverages, we also queried LSUCC0530 Pha protein sequences in PSI-BLAST against the nr database, once against the full database and once limiting the search to the SAR11 cluster. We hand-picked the following sequences from this search: 17 additional SAR11 and one *Rickettsiales* PhaC homologs, 33 additional SAR11 and 9 other *Alphaproteobacteria* species PhaA homologs, and 15 additional SAR11 PhaB homologs. We also manually added a further 14-18 homologs previously identified in a comparative genomic analysis of the SAR11 clade (7) to each protein family. All query results for each enzyme were combined into one sequence file and de-duplicated. Sequences were aligned with MUSCLE v3.8.1551 (40) and trimmed with trimAl v1.4.1 (41) using default settings. Phylogenetic trees were inferred with IQ-TREE v2.0.6 (42) using 1000 ultrafast bootstrap approximation replicates and model LG+R8 (PhaA), LG+R7 (PhaB), or LG+R9 (PhaC); visualization was performed in iTOL v6 (43).

### Plasmid construction

Primers used to construct the expression vectors in this study are listed in Table 2. Expression of *phaCBA_FU_* was achieved by ligating *phaC* (Ga0258990_1170) to *phaBA* (Ga0258990_1172‒1173) cloned from the LSUCC0530 genome and inserting the fusion into the multiple cloning site of pBBR1MCS-2 (44). Expression of *phaCAB_CN_* was achieved by inserting *phaC1AB1* (H16_A1437‒A1439) cloned from the *C. necator* H16 genome into the multiple cloning site of pBBR1MCS-2.

**Table 2.**
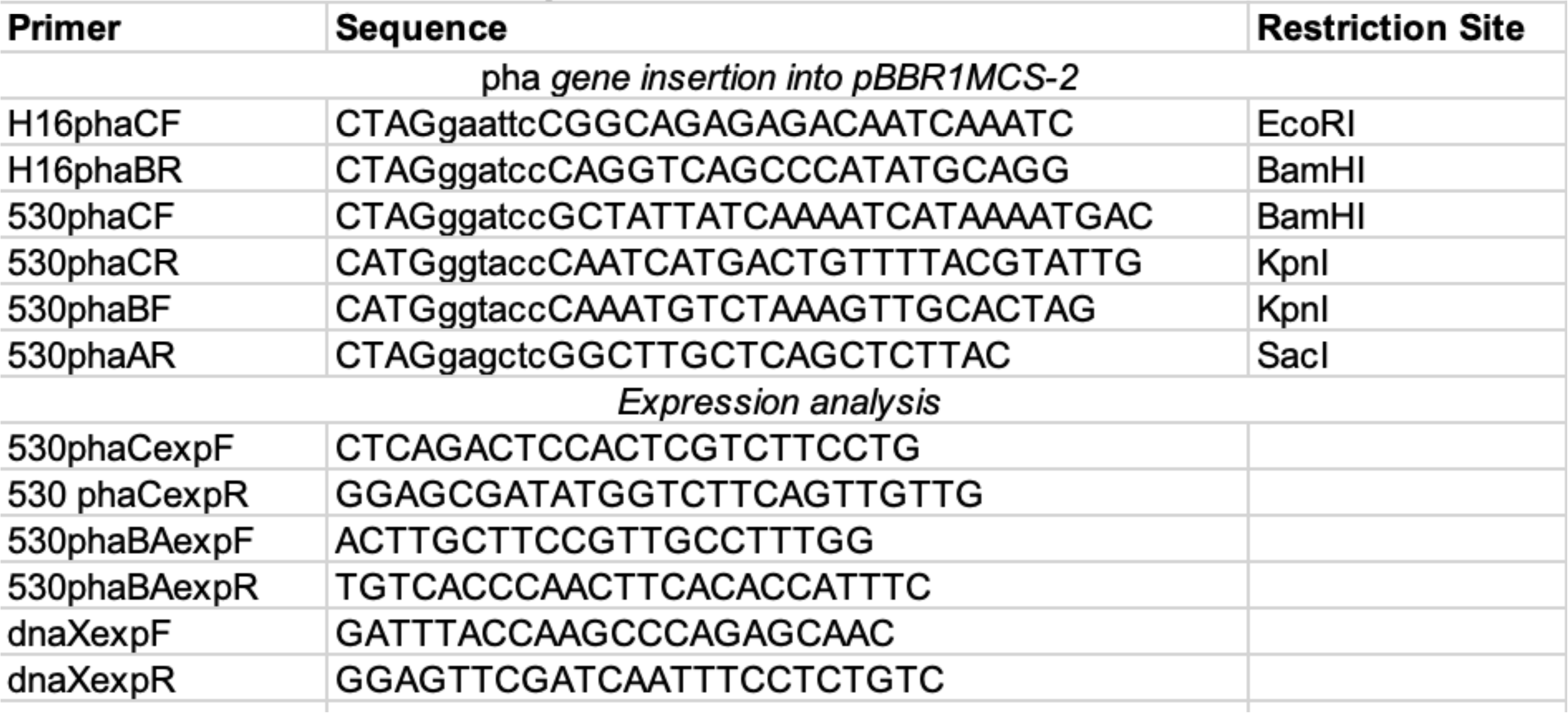
Primers used in this study.

### Lipid extraction and GC/MS

We collected LSUCC0530 cells grown in 2 liters CCM5PK or *C. necator* H16 cells grown in 50 ml nutrient broth supplemented with sodium gluconate by vacuum filtration onto 25-mm diameter, 0.1-µm pore polyvinylidene fluoride membrane filters (Durapore, Darmstadt), which were placed in a 50-ml conical polypropylene tube and frozen at −80°C. We extracted lipids from cells on filters and performed GC/MS similarly to a previously described protocol (45). Briefly, we added 1.5 ml chloroform and 1.5 ml acidified methanol to glass culture tubes containing the filters (enough to fully submerge the filters in solvent), capped the tubes with screw-tops containing PTFE/silicone septa, and vortexed for 1 min. Tubes were incubated for 2.5 h at 100°C on a heating block, then cooled on ice for 5 min. We washed samples by adding 500 µl double-deionized H2O and vortexing briefly. We removed the bottom, organic layer into 150 µl inserts in GC glass vials and capped these with PTFE/silicone septa. Samples were analyzed by GC/MS on an 7890A GC system (Agilent Technologies, Santa Clara, CA) equipped with a DB-WAX UI column as stationary phase coupled with an Agilent 5975C inert XL EI/CI Mass Selective Detector with Triple-Axis Detector. The GC temperature gradient was programmed as follows: initialization, 80°C for 2 min; ramp 1, to 210°C at 10°C/min; ramp 2, to 250°C at 50°C/min, hold 1 min. We used poly((R)-3-hydroxybutyrate-co-(R)-3-hydroxyvalerate-co-(R)-3-hydroxyhexanoate) (Sigma-Aldrich, USA) to generate the standard curve.

### Reverse transcription and quantitative PCR

We extracted RNA from biological triplicates of LSUCC0530 cultures as follows: Cells of LSUCC0530 grown in 1 liter JW5 with 1x (2 mM) or 10x (20 mM) ammonium (NH_4_Cl) and 1x (100 µM) or 10x (1 mM) phosphate (KH_2_PO_4_), as noted, were collected onto 25-mm, 0.1-µm pore polyethersulfone membrane filters. 2 ml RNAprotect Bacteria Reagent (Qiagen, Hilden) was added to each filter and incubated for 15 min at room temperature before vacuuming through the filter. Each filter was placed in a 2-ml microcentrifuge tube, to which 300 µl lysozyme (15 mg/ml in10 mM Tris-Cl and 1mM EDTA) was added, and vortexed for 10 s; filters were then shaken for 5 min to lyse cells. 1.5 ml Buffer RLT Plus (Qiagen) containing 15 µl β-mercaptoethanol and 1.125 µl Reagent DX (Qiagen) was added and tubes were vortexed vigorously. RNA was extracted from each filter with an AllPrep kit (Qiagen), including treatment with RNase-Free DNase (Qiagen) per manufacturer protocol. Extracted RNA was treated with 2 µl DNase RQ1 (Promega) for 60 min to remove any remaining contaminating DNA.

We synthesized cDNA from RNA using Luna RT Mix (New England Biolabs, Ipswich, MA) and measured gene expression by qPCR performed with 2X Universal SYBR Green Fast qPCR Mix (ABclonal, Woburn, MA) under the following conditions: 95°C for 3 min; 45 cycles of 95°C for 5 s, 57°C for 10 s, 60°C for 35s; and melting curve acquisition from 60°C to 95°C. We confirmed using serial 10-fold dilutions of PCR-amplified target DNA that the qPCR primer pairs (listed in Table 2) had efficiencies between 90 and 110% under these conditions. Cycle thresholds (Ct) for each sample were normalized to those of the reference gene *dnaX* (ΔCt = target gene Ct – *dnaX* Ct); ΔΔCt values were calculated by subtracting the average ΔCt for the 1x JW5 cultures; fold changes were calculated as 2^(−ΔΔCt).

## Results

### “*Ca*. F. ubique” replicates without a supplemental carbon source

We observed that when transferred from a carbon-replete medium into a medium lacking a supplemental carbon source, LSUCC0530 grew at nearly the same rate (minimum doubling time 2.19 ± 0.27 d and 2.02 ± 0.01 d, respectively) and approached a similar cell density (3.5E6 ± 2.8E6 cells/ml and 8.3E6 ± 1.3E6 cells/ml, respectively) as when transferred into fresh carbon-replete medium (Fig 1A). Indeed, when we serially transferred LSUCC0530 into new cultures lacking supplemental carbon, the cells continued to grow through up to nine transfers (Fig 1B; Fig S3), representing over 80 generations. This ability of LSUCC0530 to grow without a carbon source is notably different from other well-studied SAR11 strains such as “*Ca*. Pelagibacter ubique” HTCC1062, which was unable to grow upon inoculation into medium without supplemental carbon (Fig 1C).

**Fig 1.**
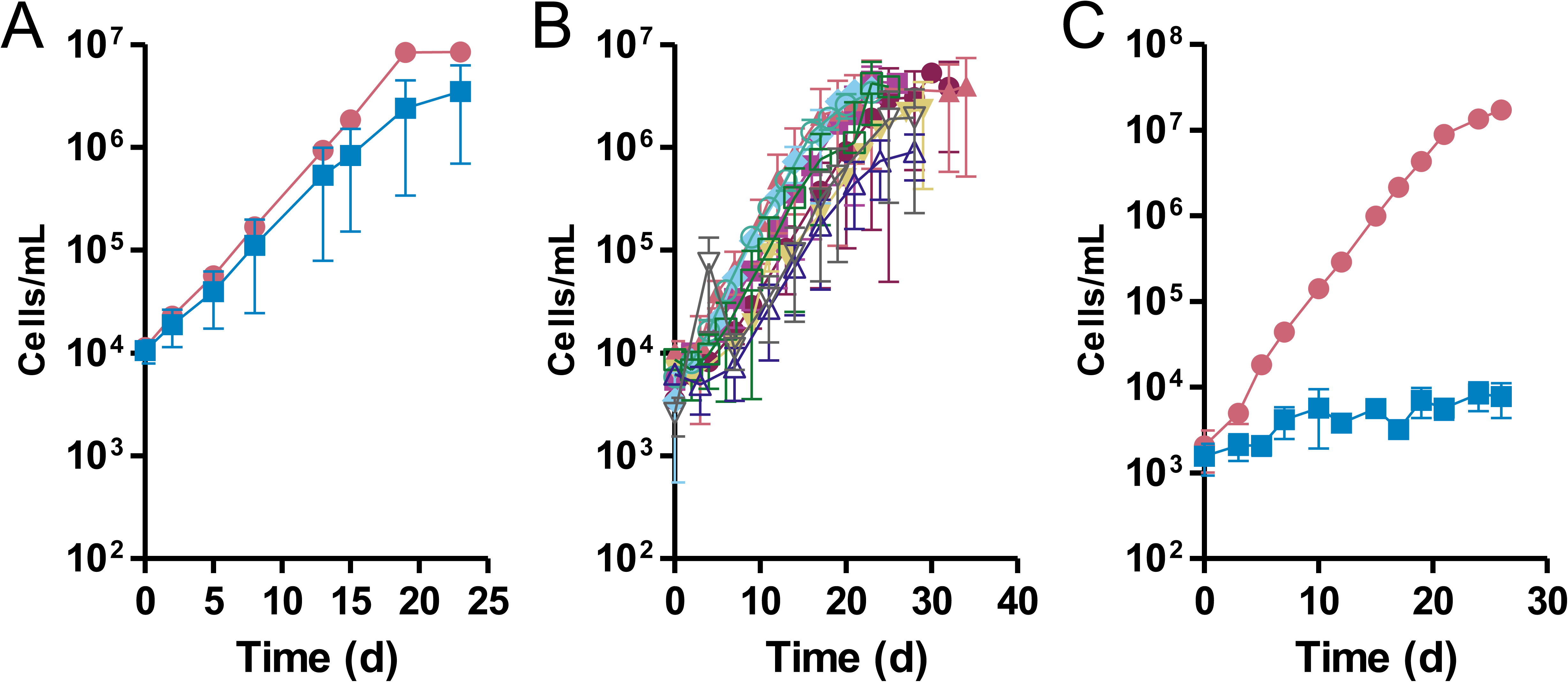
LSUCC0530 grows for many generations without a supplemental carbon source. A) The rate of growth was measured for LSUCC0530 grown in JW5 with (circles) and without (squares) supplemental carbon. B) The rate of growth was measured for LSUCC0530 grown in JW5 without supplemental carbon over nine serial transfers (T1‒9); closed circles, T1; closed squared; T2, closed triangles, T3; closed inverted triangles, T4; diamonds, T5; open circles, T6; open squares, T7; open triangles, T8; open inverted triangles, T9. C) The rate of growth was measured for “*Ca*. Pelagibacter ubique HTCC1062 grown in AMS1 with (circles) and without (squares) supplemental pyruvate. Results represent the means from three biological replicates ± 1 standard deviation.

### “*Ca.* F. ubique” cells make polar, Nile red-staining inclusion bodies

Since we had previously noted the presence of PHA production genes in the LSUCC0530 genome (34), the ability of LSUCC0530 to multiply in the absence of supplemental carbon raised the question of whether this organism could use stored carbon during starvation periods. In evaluating the morphology of LSUCC0530 cells via transmission electron microscopy (TEM), we noted that many of the cells contained a single polar, ground glass-like inclusion body (Fig 2A). With the notion that these inclusion bodies might be carbon storage granules, we stained LSUCC0530 cells with Nile red, a lipophilic dye that produces high-intensity fluorescence when bound to PHA (46). Imaging via epifluorescence microscopy demonstrated that a fraction of LSUCC0530 cells contained a single, polar inclusion body that stained brightly with Nile red (Fig 2B). These Nile red-staining inclusion bodies occurred in the same subcellular location as the inclusions we observed in the TEM images; when seen in dividing cells, the inclusions were always found at the pole opposite the site of cell division (Fig 2A). Based on analysis of the size of the cellular inclusions in the TEM images, assuming the inclusions were composed of 100% PHB, we estimated that LSUCC0530 cells contained an average of 2.93 fg/cell (1.24‒4.63 fg/cell).

**Fig 2.**
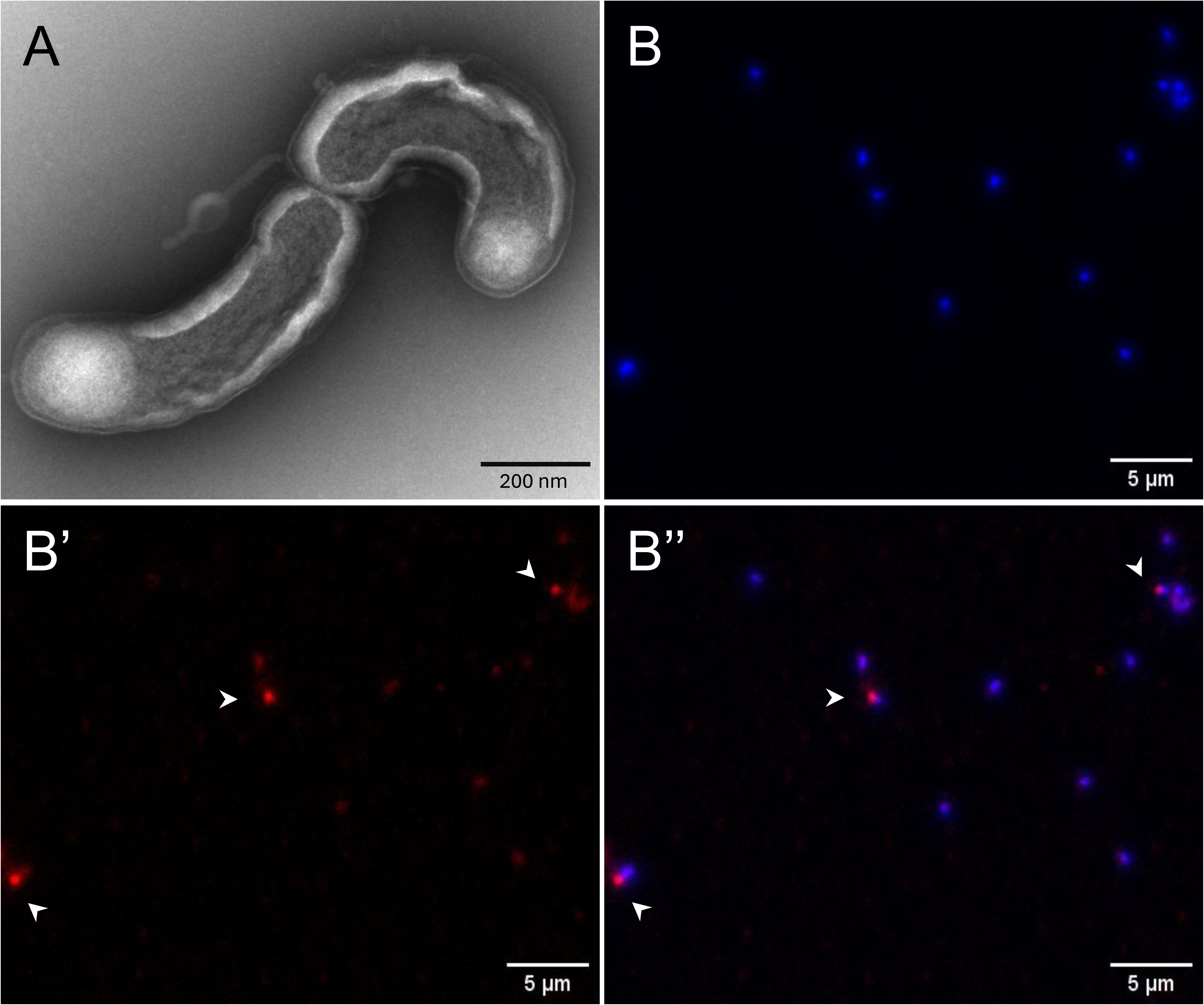
Transmission electron and fluorescence microscopy of LSUCC0530 cells. A) TEM was performed on LSUCC0530 cells grown to late exponential phase in JW5. B) LSUCC0530 cells were grown to early stationary phase in JW5, stained with Nile red (PHA stain, red) and DAPI (nucleic acid stain, blue), and imaged via epifluorescence microscopy. B, DAPI channel; B’, TRITC channel; B’’, merged image. Arrowheads indicate cells with Nile red-positive inclusions.

### The genomes of a subset of SAR11 strains encode PHA enzyme homologs

The genomic potential for PHA production was previously described in genomic characterizations of LSUCC0530 and other SAR11 strains in both subclades IIIa and LD12 (7,17). The genomes of both LSUCC0530 and the SAR11 subclade IIIa strain LSUCC0261 contain loci predicted to encode homologs of the enzymes involved in both the production of PHA (PhaC, phasin family protein PhaP, PhaB, PhaA, and short chain enoyl-CoA hydratase PhaJ; encoded in LSUCC0530 by locus tags Ga0258990_1170‒74) and its degradation (alpha/beta hydrolase PhaZ; encoded in LSUCC0530 by locus tag Ga0258990_1175). Unlike in other, well-studied organisms that produce PHA, many of which contain multiple copies of various *pha* genes (47,48), the LSUCC0530 genome contains only a single copy of the core *phaA*, *phaB*, and *phaC* genes.

Phylogenetic trees of the PhaA, PhaB, and PhaC protein sequences encoded in available SAR11 genomes demonstrated that the SAR11 homologs of these proteins formed a monophyletic group in each tree and were deeply divergent from those found in other species, including other Alphaproteobacteria (Fig S1). Our protein search found homologs for the Pha proteins throughout the subclades of SAR11, but these proteins appear to be enriched in the IIIa and LD12 subclades. Pha proteins in these two subclades clustered into monophyletic groups (Fig 3; Fig S1), however, the Pha protein trees generally did not follow the branching order of the species trees for SAR11 (Fig 3). For example, the SAR11 IIIa and LD12 PhaC clades were separated by sequences from SAR11 subclades Ia, Ic, II, and Aegean-169 sequences, even though IIIa and LD12 are taxonomically sister clades. This corroborates previous phylogenetic studies of *pha* genes and PhaC proteins that have illustrated the disconnect between 16S rRNA phylogeny and *pha* gene relatedness (18,49) and suggests intra-SAR11 horizontal transfer of *pha* genes.

**Fig 3.**
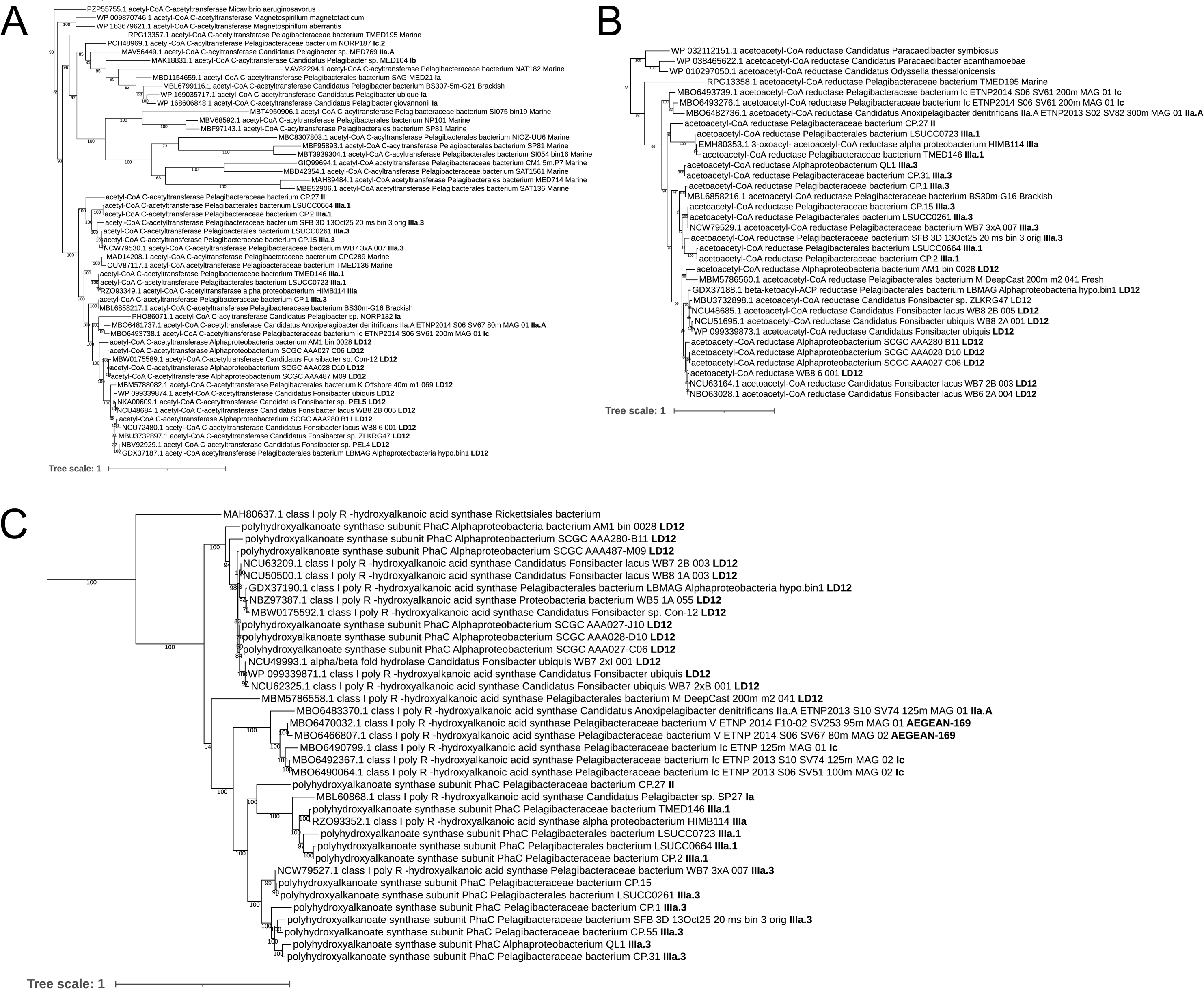
Pha protein phylogeny in the SAR11 clade. Maximum likelihood phylogenetic trees of Pha synthesis enzyme protein sequences annotated in SAR11 genomes. A) PhaA homologs; B) PhaB homologs; C) PhaC homologs. Node values indicate branch support bootstrap values (as a percentage; n=1000). Subclade assignments follow strain names; where subclade assignment is unavailable, water source type (brackish or marine) is provided. Strain names are replaced by species names for RefSeq proteins. Proteins designated FabG (B) are annotated as 3-oxoacyl- [acyl-carrier-protein] reductase, which may replace PhaB during PHA synthesis. Expanded trees containing non-SAR11 homologs are displayed in Figs S1A‒C.

### Heterologous expression of *phaCBA* from LSUCC0530 causes production of Nile red-positive inclusion bodies

To determine whether the LSUCC0530 *pha* locus is involved in making the presumptive carbon storage inclusion bodies observed in Fig 2, we heterologously expressed the LSUCC0530 homologs of the core PHA production genes *phaC*, *phaB*, and *phaA* (*phaCBA_FU_*) in *E. coli*. We performed epifluorescence microscopy on *E. coli* complemented with empty vector, vector expressing *phaCBA_FU_*, or vector expressing *phaC1AB1* from *C. necator* H16 (*phaCAB_CN_*, positive control). We observed inclusion bodies that stained with Nile red in *E. coli* strains expressing either *phaCBA_FU_* or *phaCAB_CN_* (Fig 4A and Fig 4B); no inclusions were seen in *E. coli* cells carrying empty vector (Fig 4C).

**Fig 4.**
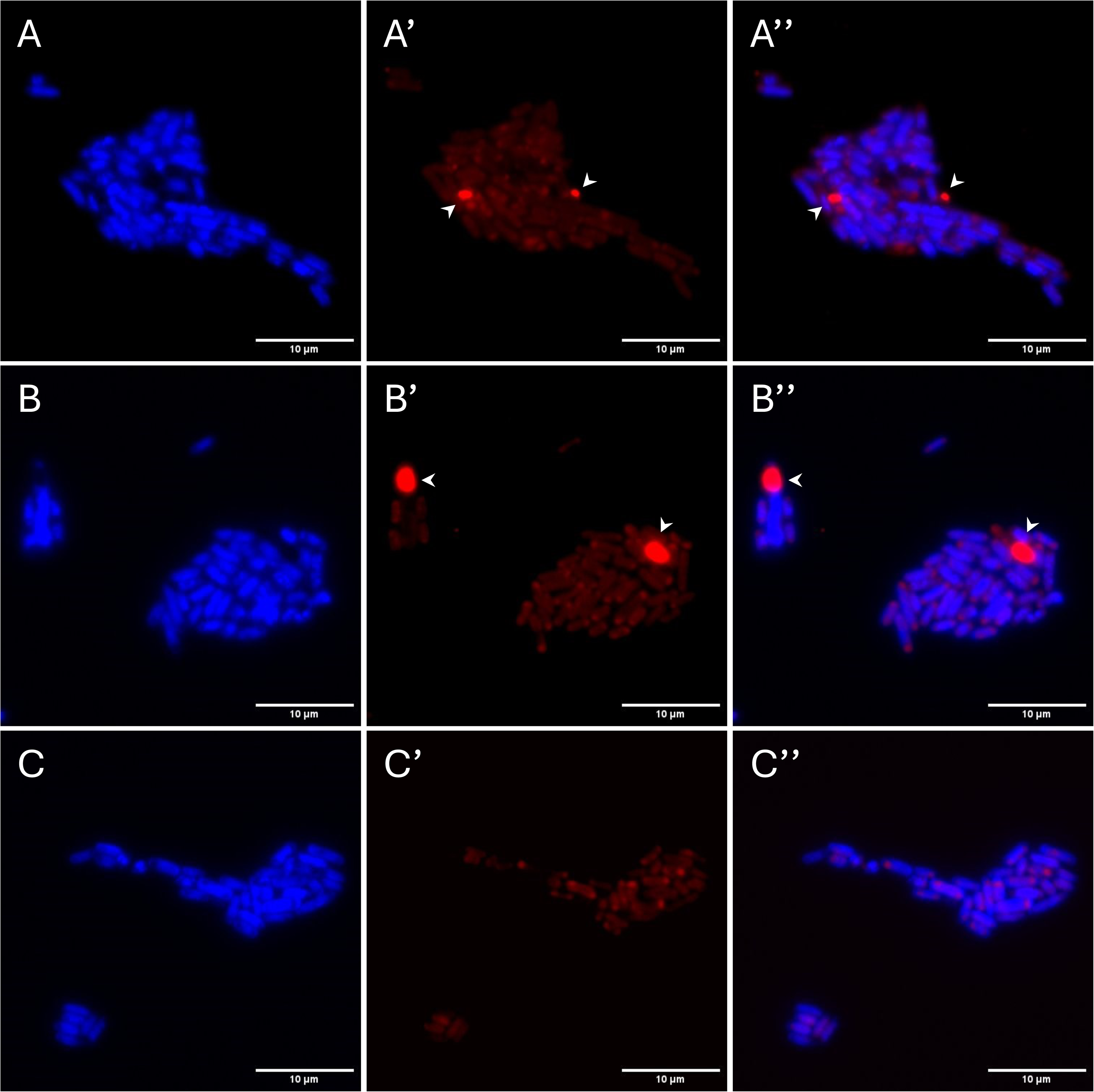
Fluorescence microscopy of *E. coli* carrying heterologous *pha* constructs. *E. coli* A) expressing *phaCBA_FU_*, B) expressing *phaCAB_CN_*, or C) carrying empty vector was stained with Nile red (PHA stain, red) and DAPI (nucleic acid stain, blue) and imaged via epifluorescence microscopy. Left, DAPI channel; middle, TRITC channel; right, merged images. Arrowheads indicate cells with Nile red-positive inclusions.

### LSUCC0530 cells produce PHB

To confirm that the inclusion bodies observed in LSUCC0530 cells (Fig 2) contained PHA, and to identify the presumptive PHA monomer(s), we conducted gas chromatography/mass spectrometry (GC/MS) on lipid extracts of LSUCC0530 cultures. The sole PHA derivative detected in the LSUCC0530 cells was 3-hydroxybuytyric acid (3HB), the monomer unit of PHB (Fig 5A). We also observed 3HB production in a *C. necator* culture grown in nutrient broth with gluconate, serving as a positive control, with a similar GC retention time to the 3HB found in LSUCC0530 (Fig 5B). No PHA monomer compounds were detected in extracts of a CCM5PK medium-only blank (Fig S2). The PHB content averaged 1.21 fg/cell (0.087‒3.39 fg/cell), which aligns with our estimate of cellular PHB content based on image analysis.

**Fig 5.**
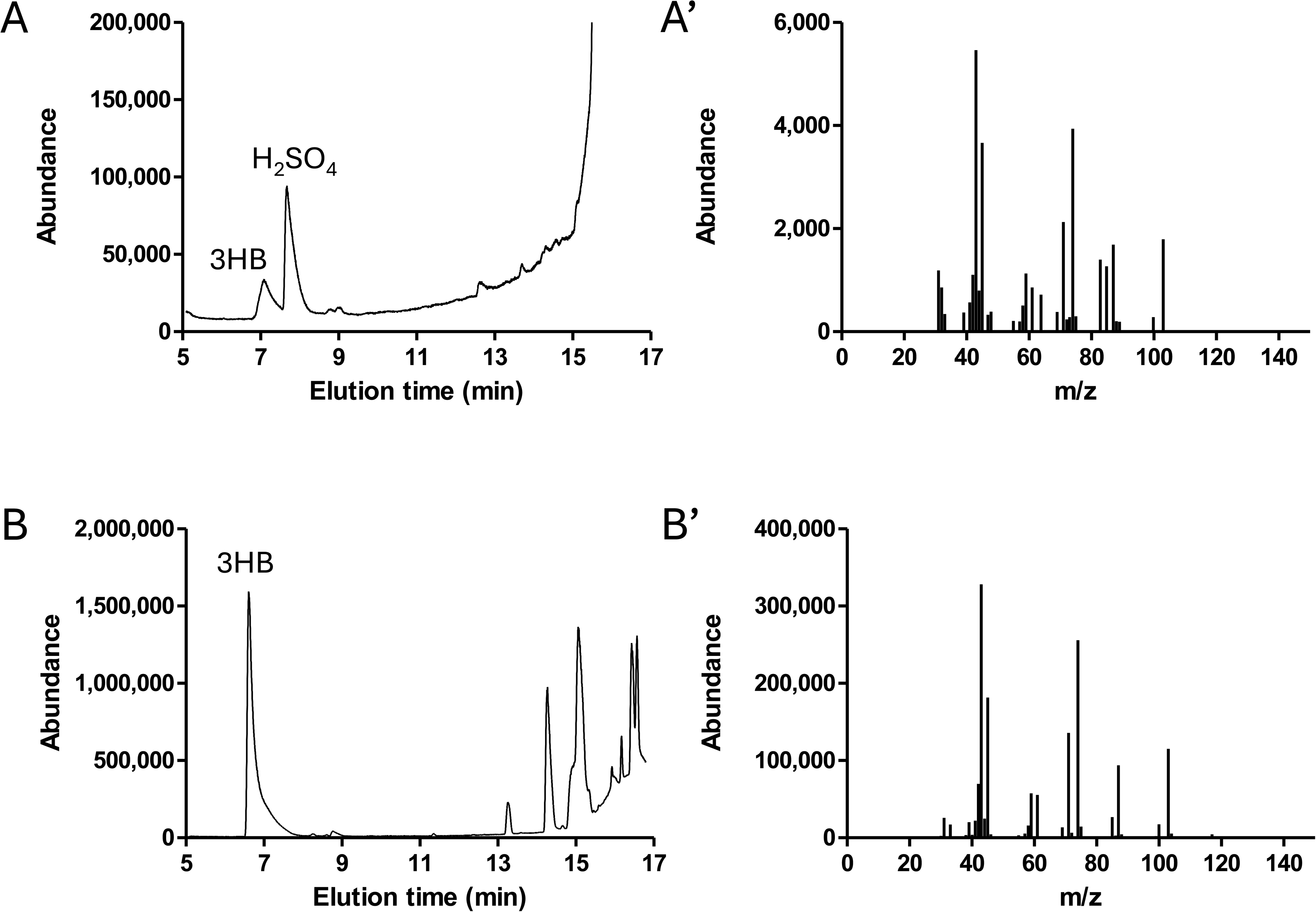
GC/MS analysis of LSUCC0530 culture lipid extracts. GC/MS was performed on lipid extracts from cultures of A) LSUCC0530 grown into early stationary phase on CCM5PK and B) *C. necator* grown into stationary phase on nutrient broth with gluconate. Left, GC chromatograms; peaks are labeled with the highest confidence match of the corresponding MS spectrum to the NIST database. Right, MS spectra for the peaks labeled 3-hydroxybutyric acid (3HB). Note the higher scale for abundance in (B). H_2_SO_4_ is a reagent used during lipid extraction.

### Nitrogen and phosphate concentrations do not affect PHA production in LSUCC0530

Nitrogen and phosphorus limitation commonly induce PHA production in many bacteria (29,30). To determine whether production of the PHA inclusions in LSUCC0530 is induced by nitrogen and/or phosphorous limitation, we quantified PHA inclusions via Nile red staining and epifluorescence imaging in LSUCC0530 cultures grown in JW5 with either the standard concentrations of ammonium and phosphate (2 mM and 100 µM, respectively) or 10x the concentrations of both compounds. We found no difference in the fraction of LSUCC0530 cells containing PHA inclusions between the two growth conditions (Fig 6A). Additionally, we wanted to know whether LSUCC0530 cells grown in a medium lacking a supplemental carbon source would consume the stored PHA and, therefore, produce fewer progeny containing PHA inclusions. There was no difference in the fraction of LSUCC0530 cells containing PHA inclusions regardless of the presence or absence of supplemental carbon in the growth medium (Fig 6A). Regardless of the growth condition, we found a high variance in the percentage of cells considered positive for PHA inclusions, from zero or near zero to greater than 40% (Fig 6A). This suggests considerable subpopulation heterogeneity that could indicate a wide range in growth phenotypes depending on how the cells use PHB.

**Fig 6.**
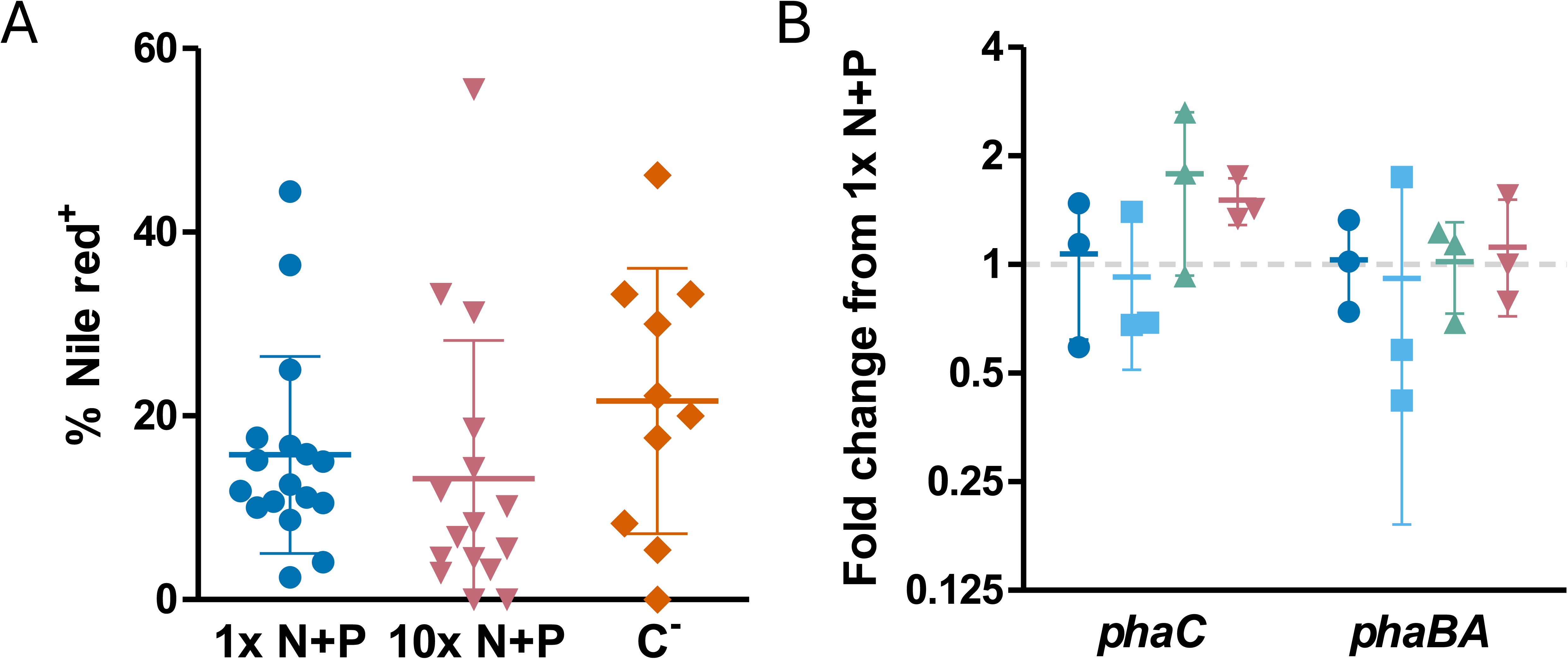
Microscopic and transcriptional analysis of nitrogen and phosphorus influence on PHA production in LSUCC0530. A) The percentage of cells containing PHA inclusion bodies was counted for LSUCC0530 cultures grown to early stationary phase in JW5 with 1x ammonium and phosphate (circles), 10x ammonium and phosphate (inverted triangles), or without supplemental carbon (diamonds) and then stained with Nile red and DAPI and imaged via epifluorescence microscopy. Each point represents one biological replicate; five fields of view were analyzed for each replicate. Error bars represent ± 1 standard deviation. B) Expression of *phaC* and *phaBA* was analyzed via RT-qPCR for cultures of LSUCC0530 grown to early stationary phase in JW5 with 1x ammonium and phosphate (circles), 10x ammonium (squares), 10x phosphate (triangles), or 10x ammonium and phosphate (inverted triangles). Each point represents one biological replicate; values displayed are the fold change from that of the average of 1x JW5. Error bars represent ±1 standard deviation.

We also investigated whether nitrogen and/or phosphorous concentrations influence expression of PHA production genes using reverse transcription-quantitative PCR (RT-qPCR) on LSUCC0530 grown in JW5 with 1x or 10x ammonium and/or phosphate. We observed no difference in the expression levels of *phaC* or *phaBA* in LSUCC0530 grown in any of the tested media (Fig 6B), suggesting that either PHB production was constitutive or that expression is responsive to different stimuli than for other known PHB-producing taxa.

## Discussion

While the genomic potential for PHA production by LSUCC0530 and other SAR11 strains has previously been identified (7,16), the activity of *pha* homolog genes had not been studied in any SAR11 strain before this work. In this study, we determined that the *pha* synthesis genes encoded in the LD12 strain LSUCC0530 were functional and that LSUCC0530 produced PHB irrespective of carbon, nitrogen, or phosphate concentration. Without tools available to modify the genomes of SAR11 bacteria, we were unable to inactivate the *pha* locus and investigate whether PHA production contributes to survival of LSUCC0530 under starvation conditions (Fig 1A). Our estimate of cellular PHB concentration at a maximum of ∼3 fg/cell, combined with the total carbon content of the similarly sized HTCC1062 at ∼6.5 fg/cell (50), meant that PHB alone could not be responsible for the continued growth of LSUCC0530 without added carbon. However, as the LSUCC0530 genome is not predicted to encode the enzymes necessary for CO_2_ fixation, we think it possible that some of this growth is achieved by cells utilizing carbon stored as PHB. We have previously reported a similar phenotype in the OM252 Gammaproteobacteria representative strain LSUCC0096 (51). Although LSUCC0096 has genes for autotrophy, it also has putative *pha* genes and can grow for multiple transfers without an added electron donor, implicating PHA storage in energy generation and possibly as a carbon supply.

The potential utilization of stored carbon by LSUCC0530 to support survival under starvation conditions may complement the function provided by proteorhodopsin, another survival mechanism widely possessed by bacteria throughout the *Pelagibacterales* (7,34,52,53). Proteorhodopsin absorbs light and pumps protons, serving to maintain membrane potential (52,54). In the subclade Ia strain HTCC1062, proteorhodopsin prevents oxidation of biomass during periods of starvation (55). Given the ability of LSUCC0530 to continue growing under starvation conditions, PHA storage capability may provide LD12, subclade IIIa, and other PHA-producing SAR11 strains with a fitness advantage during periods of starvation, since PHA oxidation may provide energy as well as serve as a carbon source. The relative contribution of proteorhodopsin to the energy needs of PHA-oxidizing SAR11 cells deserves further study to determine whether and how these metabolisms are used in conjunction.

Unlike in many other organisms, nitrogen and phosphorous concentrations do not appear to influence PHA production in LSUCC0530 (Fig 6). It is possible that PHA production in SAR11 bacteria is regulated via some other mechanism that we did not investigate. Indeed, as SAR11 are oligotrophic organisms, they are already well-adapted to the canonical nitrogen and phosphorous limitation that typically promote PHA production in other organisms (4). Less commonly observed PHA regulatory mechanisms include oxygen restriction (56), iron limitation (57), and quorum signaling (58); however, our cultures were grown aerobically with excess iron, and no evidence of quorum signaling in SAR11 bacteria has been discovered as yet (59). Transcriptional regulators shown to affect *pha* gene expression and PHA production in other bacteria include the alternative sigma factors RpoN (60) and RpoS (61), though there are no genes encoding homologs of those sigma factors annotated in the LSUCC0530 genome. Currently there are no obvious potential mechanisms for regulating PHA production in LSUCC0530, making it likely that its *pha* synthesis genes are constitutively expressed.

A possible explanation for the apparent lack of PHA synthesis regulation in LSUCC0530 may be related to the slow growth phenotype of SAR11 bacteria. The highly streamlined genomes of SAR11 bacteria contain a single copy of the 5S, 16S, and 23S rRNA genes (9), and SAR11 cells have low ratios of rRNA/rDNA, both of which are linked to slow growth rates (62,63). However, SAR11 bacteria are at least as metabolically active as other prokaryotes (64). It may be that the growth-limiting factor in SAR11 cells is the translation machinery, causing biomass production to lag behind catabolism and producing excess intracellular NADH and ATP—the very conditions that cause cells to shunt acetyl-CoA away from the TCA cycle and into PHA production (31,65). Thus, PHA production may be constitutive in those SAR11 strains whose genomes contain complete *pha* loci simply due to their particular oligotrophic physiology. In these bacteria, it is possible that the major factor affecting the presence of PHA in a cell is the rate of PHA utilization, rather than synthesis.

Surprisingly, the ratio of LSUCC0530 cells containing PHA inclusions was similar regardless of whether the bacteria were grown in media with or without a supplemental carbon source; in fact, the average proportion of PHA-containing cells was slightly higher in the carbon-deplete medium, though this was not statistically significant (*p*=0.28) (Fig 6A). It is theoretically possible that the depolymerase PhaZ encoded in the LSUCC0530 genome is nonfunctional, as we did not test this hypothesis. However, we think it more likely that PhaZ is functional and is used to depolymerize PHA during periods of carbon limitation. We speculate that those cells that have used up their PHA stores are more likely to have died off and no longer be detectable via microscopy, leaving those still with available PHA to continue growing. We note that the carbon-starved cultures used for microscopy grew to lower average densities (4.8E6 ± 3.9E6 cells/ml) than the carbon-replete cultures (1.2E7 ± 6.5E6 cells/ml), so while the ratio of Nile red-positive cells in the carbon-starved cultures was equal to or higher than that in the carbon-replete cultures, the total number of live cells was lower. It may be worthwhile in a future study to track the ratio of PHA-containing LSUCC0530 cells in carbon-starved cultures throughout a growth curve into death phase as well as over multiple transfers until cell division finally ends, to see if this ratio increases and then eventually decreases over time.

The wide range of LSUCC0530 cells containing visible PHA granules under the conditions we evaluated needs to be reconciled with our hypothesis of constitutive PHB synthesis. One possible explanation for this discrepancy is that in some cells, there is simply not enough accumulated PHA for detection. It was difficult to visualize subcellular structures in these ultramicrobacteria (2) using fluorescence microscopy, and it may be that smaller amounts of PHB in these cells are indistinguishable from background Nile red staining of the cell membrane. Additionally, the relative amount of PHB in each cell may reflect different amounts of utilization stemming from phenotypic heterogeneity among individual cells. If PHB production is constitutive and the amount found in a cell is due to the relative rate of consumption vs. production, then the amount of PHB may reflect higher or lower rates of a cell’s metabolic activity. For example, by modeling viability of LD12 in natural populations, we estimated that over half of LD12 cells may be dormant at any one time (66). If similar levels of quiescence occur in culture, that may partially explain the large variance in cellular PHB levels.

Genes encoding PHA enzyme homologs are found across all the major SAR11 subclades (Fig 3, Fig S1). These genes have previously been reported in the brackish and freshwater SAR11 subclades II, IIIa, and LD12 (7,16,17), but to our knowledge this is the first work to point out the presence of these genes within marine-dwelling SAR11 subclade I and Aegean-169. Though the potential for PHA production can be found across the entire SAR11 clade, only a subset of available SAR11 genomes encode homologs of Pha enzymes. The *pha* genes generally grouped by subclade (Fig 3), although the subclade branching order was different from that of species trees generated with whole genomes (7). Furthermore, *pha* genes were much more common in the fresh and brackish-water subclades LD12 and IIIa compared to other SAR11 subclades. Coupling these results with the observation that SAR11 Pha proteins are deeply divergent from those found in other Alphaproteobacteria suggests that the genes enabling PHA production were inherited by a common ancestor of the IIIa and LD12 lineages and then horizontally transferred very occasionally to other lineages within SAR11.

This study is the first to examine PHA production by a member of the *Pelagibacterales*. Pressing questions remain regarding the heterogeneity of PHA accumulation in LSUCC0530 cells under all tested conditions *in vitro*, and whether PHA granules would be seen in similar fractions of LSUCC0530 cells taken from freshwater environments undergoing either eutrophication or oligotrophication. It would also be worthwhile to explore whether the putative *pha* homologs present in cultivated SAR11 strains from the marine- and brackish-dwelling subclades are active and if these strains can continue propagating in the absence of a carbon source, as with the freshwater-evolved LSUCC0530. Further work should also investigate the possibility that other likely PHA-producing SAR11 strains have alternative regulatory mechanisms governing PHA production, and if so whether these mechanisms also exist in LSUCC0530.

## Acknowledgements

We would like to thank Dr. Moh El-Naggar and Dr. Tingting Yang at University of Southern California for access to and training on fluorescence microscopy. We thank Dr. Hagen Buck-Wiese at USC for critical review of our analyses regarding LSUCC0530 PHB utilization for cellular growth. We would also like to thank Dr. Ying Xiao in the Shared Instrumentation Facility at Louisiana State University for help with transmission electron microscopy. We thank Chuankai Cheng at the University of Southern California for providing the CCM5 recipe. We acknowledge the computational resources provided by the Center for Advanced Research Computing at the University of Southern California. This work was supported by the Simons Foundation Early Career Investigator in Marine Microbial Ecology and Evolution Award to JCT.

## Data availability

The datasets and images generated in the work presented here are available from the corresponding author upon reasonable request.

## Supplementary Information

**Fig S1.**
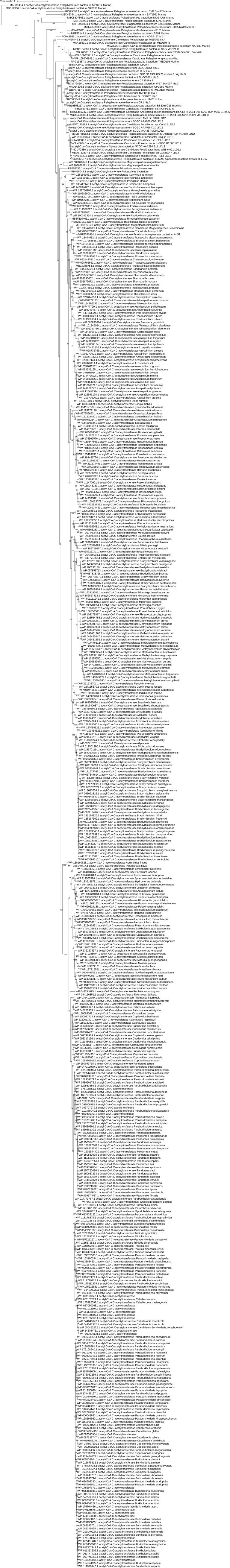
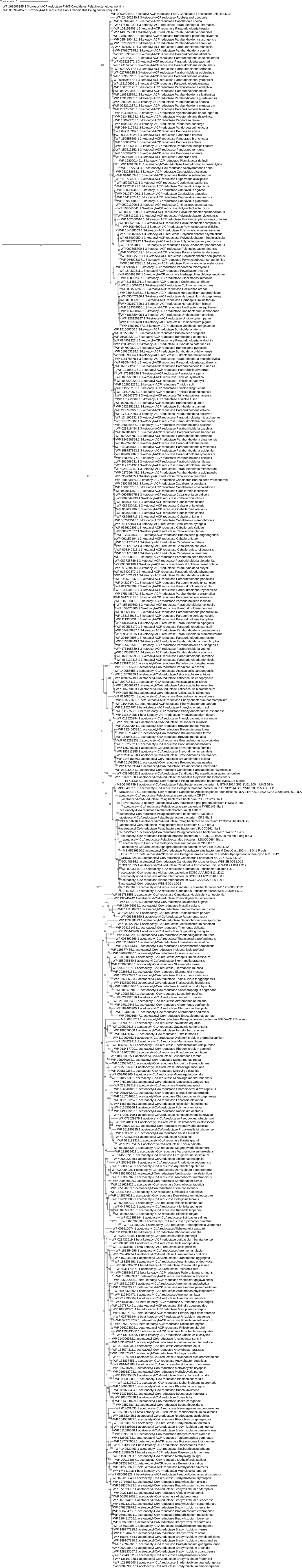
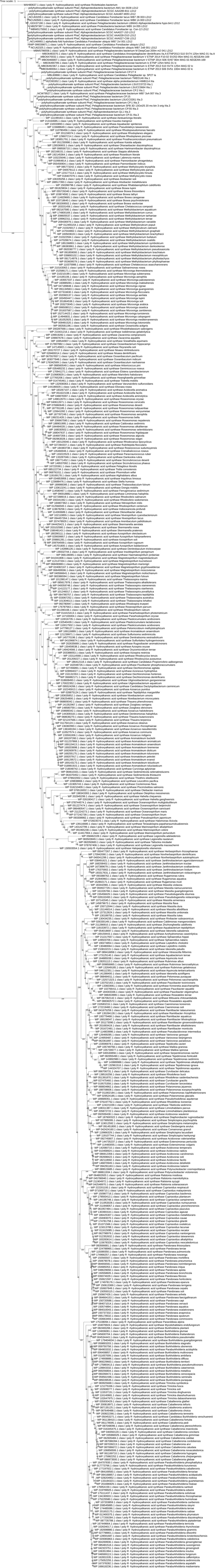
Expanded Pha protein phylogeny. Maximum likelihood phylogenetic trees of Pha synthesis enzyme protein sequences annotated in bacterial genomes. A) PhaA homologs; B) PhaB homologs; C) PhaC homologs. Node values indicate branch support bootstrap values (max=100). SAR11 subclade assignments follow strain names; where subclade assignment is unavailable, water source type (fresh, brackish, or marine) is provided. Strain names are replaced by species names for RefSeq proteins. Proteins designated FabG (B) are annotated as 3-oxoacyl-[acyl-carrier-protein] reductase, which may replace PhaB during PHA synthesis.

**Fig S2.**
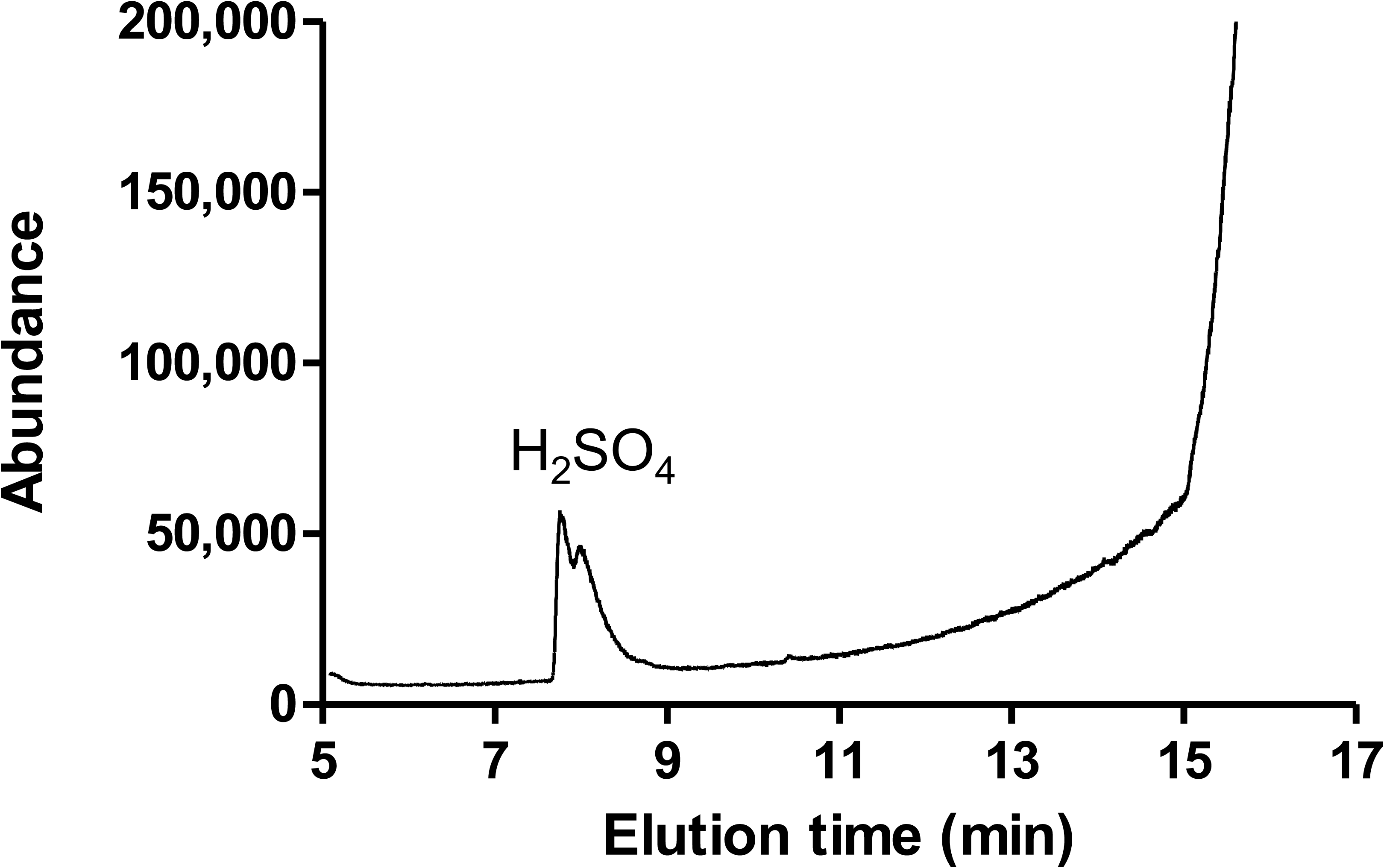
GC/MS analysis of medium blank lipid extract. GC/MS was performed on a lipid extract from a CCM5PK medium blank. GC chromatogram is shown; the peak is labeled with the highest confidence match of the corresponding MS spectrum to the NIST database. H_2_SO_4_ is a reagent used during lipid extraction.

**Fig S3.**
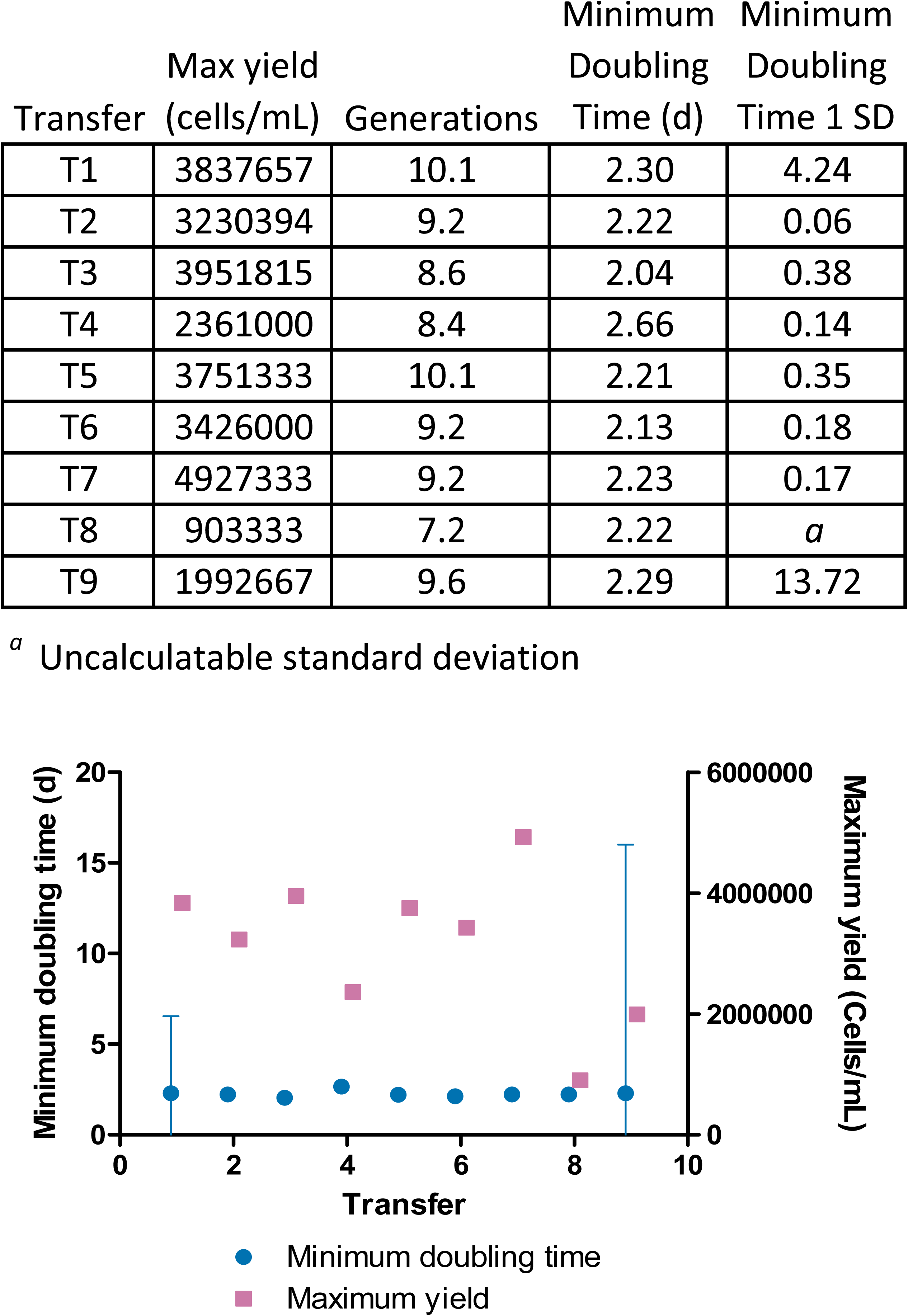
Growth rate, yield, and generation calculations for carbon-limited growth curves. Maximum growth yield, number of generations, and minimum doubling times were calculated for each LSUCC0530 carbon-limited culture depicted in Fig 1B. Circles, minimum doubling time; squares, maximum growth yield.

**Table S1.** Media used in this study.

| JW5 |  | CCM5PK |  |
| --- | --- | --- | --- |
| Component | Concentration | Component | Concentration |
| *Salts and buffer* |  | *Salts and buffer* |  |
| NaCl | 16.9 mM | NaCl | 16.9 mM |
| KCl | 416 μM | KCl | 416 μM |
| NaHCO<sub>3</sub> | 10 mM | NaHCO<sub>3</sub> | 10 mM |
| Na<sub>2</sub>SO<sub>4</sub> | 1.25 mM | MgCl<sub>2</sub>·6H<sub>2</sub>O | 2.19 mM |
| NaBr | 33 μM | CaCl<sub>2</sub>·2H<sub>2</sub>O | 434 μM |
| H<sub>3</sub>BO<sub>3</sub> | 16.2 μM | KH<sub>2</sub>PO<sub>4</sub> | 100 μM |
| SrCl<sub>2</sub> | 3.78 μM | *Transition metals* | |
| NaF | 3.1 μM | FeSO<sub>4</sub>·7H<sub>2</sub>O | 101 nM |
| KH<sub>2</sub>PO<sub>4</sub> | 6.47 μM | Nitrilotriacetic acid, disodium salt | 345 nM |
| MgCl<sub>2</sub>·6H<sub>2</sub>O | 2.19 mM | MnCl<sub>2</sub>·4H<sub>2</sub>O | 9.1 nM |
| CaCl<sub>2</sub>·2H<sub>2</sub>O | 434 μM | ZnSO<sub>4</sub>·H<sub>2</sub>O | 1.11 nM |
| *Transition metals* |  | CoCl<sub>2</sub> | 500 pM |
| FeSO<sub>4</sub>·7H<sub>2</sub>O | 101 nM | Na<sub>2</sub>MoO<sub>4</sub> | 319 pM |
| Nitrilotriacetic acid, disodium salt | 345 nM | Na<sub>2</sub>SeO<sub>3</sub> | 1 nM |
| MnCl<sub>2</sub>·4H<sub>2</sub>O | 9.1 nM | NiCl<sub>2</sub> | 1 nM |
| ZnSO<sub>4</sub>·H<sub>2</sub>O | 1.11 nM | *Vitamins* |  |
| CoCl<sub>2</sub> | 500 pM | Thiamine | 10.02 μM |
| Na<sub>2</sub>MoO<sub>4</sub> | 319 pM | Riboflavin | 13.82 nM |
| Na<sub>2</sub>SeO<sub>3</sub> | 1 nM | Niacin | 16 μM |
| NiCl<sub>2</sub> | 1 nM | Pantothenic acid | 8.5 μM |
| *Vitamins* | | Pyridoxine | 10 μM |
| Thiamine | 501 nM | Biotin | 80.2 nM |
| Riboflavin | 691 pM | Folic acid | 80.2 nM |
| Niacin | 800 nM | Cobalamin | 14.02 nM |
| Pantothenic acid | 425 nM | Myo-inositol | 10 μM |
| Pyridoxine | 500 nM | 4-Aminobenzoic acid | 1.2 μM |
| Biotin | 4.01 nM | *Carbon and nitrogen sources* |  |
| Folic acid | 4.01 nM | NH<sub>4</sub>Cl | 2 mM |
| Cobalamin | 701 pM | Sodium pyruvate | 999.6 μM |
| Myo-inositol | 500 nM | α-Ketoglutaric acid | 691 μM |
| 4-Aminobenzoic acid | 60 nM | L-Methionine | 20.1 μM |
| *Carbon and nitrogen sources* |  |  |  |
| MEM Amino Acids (50x) Solution (Sigma-Aldrich) | 0.001x |  |  |
| L-Glutamine | 500 nM |  |  |
| Dextrose | 500 nM |  |  |
| D-Ribose | 500 nM |  |  |
| Sodium pyruvate | 500 nM |  |  |
| Sodium citrate | 500 nM |  |  |
| Oxaloacetic acid | 500 nM |  |  |
| Sodium acetate | 500 nM |  |  |
| Sodium succinate | 500 nM |  |  |
| α-Ketoglutaric acid | 500 nM | | |
| Urea | 5 μM | | |
| Octanoic acid | 500 nM |  |  |
| Decanoic acid | 500 nM |  |  |
| Isobutyric acid | 500 nM |  |  |
| Butyric acid | 500 nM |  |  |
| Valeric acid | 500 nM |  |  |
| NaNO<sub>3</sub> | 38 μM | | |
| NaNO<sub>2</sub> | 2 μM | | |
| NH<sub>4</sub>Cl | 5 μM | | |

